# A multi-modal cell-free RNA language model for liquid biopsy applications

**DOI:** 10.1101/2025.03.26.645557

**Authors:** Mehran Karimzadeh, Aiden M. Sababi, Amir Momen-Roknabadi, Nae-Chyun Chen, Taylor B. Cavazos, Sukh Sekhon, Jieyang Wang, Rose Hanna, Alice Huang, Dang Nguyen, Selina Chen, Ti Lam, Kimberly H. Chau, Anna Hartwig, Lisa Fish, Helen Li, Babak Behsaz, Fereydoun Hormozdiari, Babak Alipanahi, Hani Goodarzi

## Abstract

Cell-free RNA (cfRNA) profiling has emerged as a powerful tool for non-invasive disease detection, but its application is limited by data sparsity and complexity, especially in settings with constrained sample availability. We introduce Exai-1, a multi-modal, transformer-based generative foundation model that integrates RNA sequence embeddings with cfRNA abundance data to capture biologically meaningful representations of circulating RNAs. By leveraging both sequence and expression modalities, Exai-1 captures a biologically meaningful latent structure of cfRNA profiles. Pre-trained on over 306 billion tokens from 8,339 samples, Exai-1 enhances signal fidelity, reduces technical noise, and improves disease detection by generating synthetic cfRNA profiles. We show that self-attention and variational inference are particularly important for preservation of biological signals and contextual relationships. Additionally, Exai-1 facilitates cross-biofluid translation and assay compatibility through disentangling biological signals from confounders. By uniting sequence-informed embeddings with cfRNA expression patterns, Exai-1 establishes a transfer learning foundation for liquid biopsy, offering a scalable and adaptable framework for next-generation cfRNA-based diagnostics.

## Introduction

Blood surveillance technologies have advanced significantly, particularly in cancer detection, shifting from traditional tissue-based diagnostics to less invasive liquid biopsies. For example, in cancer, liquid biopsies analyze tumor biomarkers in biofluids to detect disease, leveraging circulating tumor DNA (ctDNA), circulating tumor cells (CTCs), cell-free RNAs (cfRNAs), and other circulating biomolecules (Crowley et al., 2013; Corcoran and Chabner, 2018; Millner et al., 2013; Roskams-Hieter et al., 2022; Sundling and Lowe, 2019; Cescon et al., 2020). These approaches, in addition to applications for early detection, have also become powerful tools for minimal residual disease detection and treatment monitoring (Ignatiadis et al., 2021).

We previously identified a novel class of small RNAs (smRNAs) that are actively expressed in cancer cells but largely absent in healthy cells (Fish et al., 2018). A significant fraction of these cancer-emergent RNAs, which we named orphan non-coding RNAs (oncRNAs), are actively secreted by tumors into circulation at levels higher than ctDNA, which is mostly shed through cell death, making them strong indicators of cancer cell states. We have shown that their detection in blood opens new avenues for early cancer diagnosis, minimal residual disease monitoring, and cancer subtype stratification from small blood volumes (Fish et al., 2018; Wang et al., 2024; Karimzadeh et al., 2024).

A major challenge is detecting individual oncRNAs in small blood volumes due to the sparsity of tumor-released molecules, particularly in early-stage disease. However, circulating oncRNAs do not appear in isolation but rather in distinct patterns and profiles—akin to a constellation of stars in a starry sky. This is precisely the type of problem where AI excels. To harness the diagnostic potential of oncRNAs, we developed Orion, a deep generative model that effectively maps cfRNA fingerprints in blood to disease presence and outcomes (Karimzadeh et al., 2024, 2023). Training such models requires large datasets—often hundreds or thousands of samples—typically feasible only for common cancers. Detecting rare cancers, therefore, necessitates foundational models capable of generalization and few-shot learning, effectively recognizing disease patterns from a limited number of instances.

In recent years, large masked language models have emerged as capable foundation models, serving as powerful tools for zero-shot and few-shot learning. Models such as GPT-4 (OpenAI, 2023) leverage masked language modeling on a large number of training tokens to learn complex, context-dependent feature representations across diverse domains. This paradigm shift has begun reshaping genomics (Consens et al., 2025), where transformer-based architectures now play a key role in uncovering regulatory patterns in DNA and RNA sequences.

Several large genomic foundation models have emerged, each tailored to specific aspects of molecular biology. DNABERT (Ji et al., 2021), Borzoi (Linder et al., 2023) and GET (Fu et al., 2023) focus on linking genetic sequences to regulatory functions. Grover (Sanabria et al., 2023), a DNA foundation model, employs byte-pair tokenization to better capture genomic sequence structure. Evo-2, a long-context transformer pre-trained on all genomes, demonstrates strong performance in tasks such as gene essentiality prediction and synthetic sequence generation (Brixi et al., 2025). BigRNA (Celaj et al., 2023), specifically trained on RNA-seq datasets, excels at predicting tissue-specific RNA expression.

Single-cell foundation models have also made significant strides in transcriptomics. Models such as scBERT (Yang et al., 2022), Exceiver (Connell et al., 2022), GeneCompass (Yang et al., 2023), scFoundation (Hao et al., 2023), Geneformer (Theodoris et al., 2023), scGPT (Cui et al., 2024), and UCE (Rosen et al., 2023) have demonstrated efficacy in tasks such as cell type classification and gene expression imputation. While these models have shown promise, their full potential in genomics is still being realized (Kedzierska et al., 2023; Boiarsky et al., 2023; Liu et al., 2023).

Despite these recent advances in transformer architectures, their application to liquid biopsy data remains largely unexplored, primarily due to the limited availability of large-scale datasets. One major challenge in cfRNA analysis is the imbalance between the high dimensionality of circulating RNA features and the small number of available clinical samples. cfRNA encompasses the entirety of circulating RNA species, the majority of which are *<* 200 nt and are released in extracellular vesicles and lipoprotein complexes (Larson et al., 2021). Traditional predictive models typically focus on specific supervised tasks with a restricted feature space, often failing to capture the complex interactions between RNA species originating from different cell types. This limitation stems from the scarcity of labeled data, making it difficult to develop models that generalize across diverse biological contexts.

Although cfRNA analysis presents unique challenges, there are inherent biological structures that can be leveraged to overcome them. Importantly, cfRNAs are not independent entities, but rather share information between their sequence and expression (Wang et al., 2024; Zirak et al., 2024), e.g., those that are generated from the same regulatory pathways. Their biotypes and secretion patterns are dictated by their sequence and structure, while their expression levels are shaped by cell-type specificity and regulatory mechanisms. Because of this interplay, circulating RNAs do not appear in isolation but rather in distinct patterns. As a result, unlike conventional biomarkers, which are often interpreted deterministically, cfRNA detection must be approached probabilistically, taking into account the broader context of other RNAs within the same sample. When viewed through this multi-modal lens, the issue of data sparsity becomes more tractable.

More importantly, we have also developed an automated cfRNA assay capable of processing hundreds of liquid biopsy samples per batch, ensuring consistency and scalability. This expansive dataset provides a unique opportunity to train transformer-based masked language models, which excel at learning contextual dependencies through self-supervised learning. Here, we introduce Exai-1, a multi-modal transformer-based generative model designed specifically for cell-free small RNA data, trained and evaluated on over 13,000 plasma and serum samples. By integrating sequence and structure-informed embeddings from RNA-FM alongside cfRNA abundance data, Exai-1 unites two complementary modalities into a unified latent space, enabling it to extract biologically meaningful relationships between cfRNA sequences and their expression patterns. This multi-modal framework allows Exai-1 to generalize across diverse downstream tasks, improving signal fidelity, mitigating noise, and enhancing the ability to distinguish disease-associated cfRNA profiles. By incorporating sequence-driven knowledge with cfRNA abundance, Exai-1 provides a scalable and unified framework for understanding cfRNA biology and advancing liquid biopsy applications.

## Results

### Exai-1: training a multi-modal foundation model on large-scale cfRNA data

To train a foundational model that is unbiased toward specific patient cohorts, it is essential to select features representing broad blood biology rather than any particular pathology. We therefore curated a comprehensive feature set from our in-house annotated collection of cell free RNA (cfRNA)s, comprising 9,491,734 features. Our selection focused specifically on tRNA, snoRNA, miRNA, yRNA, lncRNA, and oncRNA biotypes. First, we retained only those cfRNAs expressed in at least 1% of samples and subsequently applied a coefficient-of-variation cutoff to preserve highly variable features (Supplementary Figure 1a–e). Given the large initial number of oncRNAs (394,175) after this selection, we used embeddings from a recently published RNA foundation model (RNA-FM) (Shen et al., 2024) to both cluster these oncRNAs and provide sequence-informed initialization for Exai-1. Specifically, we extracted the top 32 principal components (PCs) from the RNA-FM embedding space—collectively explaining over 91% of the variation, and used these PCs to initialize Exai-1’s feature embeddings for cfRNA features. We then selected 4,558 representative oncRNA features evenly distributed across this low-dimensional space (Methods; (Supplementary Figure 1e–f)). The final feature set consisted of 704 tRNAs, 610 snoRNAs, 761 yRNAs, 716 miRNAs, and 4,558 oncRNAs, each forming distinct clusters in the RNA-FM-based embedding space (Figure 1a).

**Figure 1:**
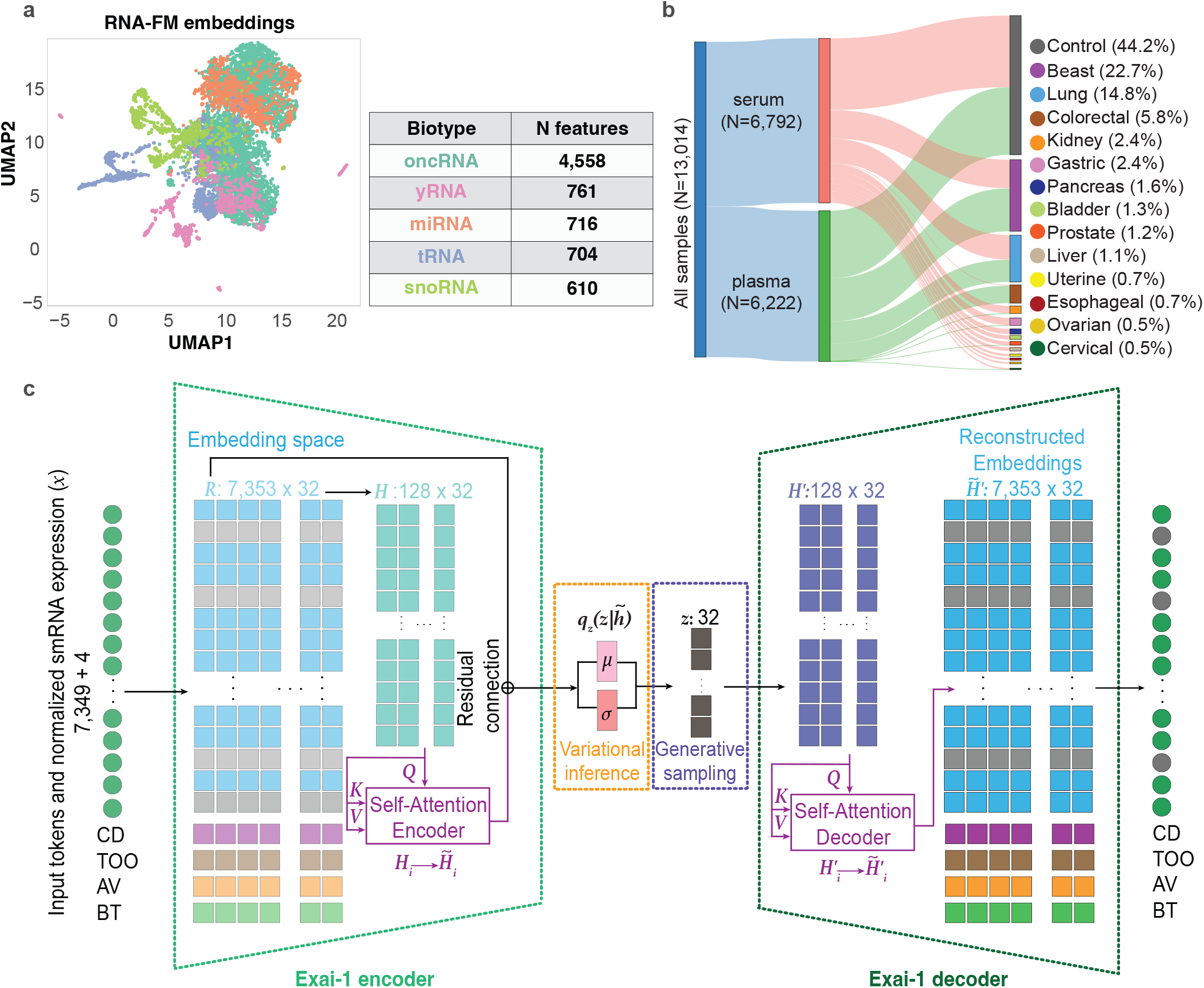
Feature selection, Data composition, and Exai-1 architecture. **(a)** Uniform Manifold Approximation and Projection (UMAP) projection of the 7,349 selected cfRNA features, color-coded by biotype (miRNA, tRNA, snoRNA, oncRNA, yRNA), showing distinct clustering patterns. **(b)** Sankey diagram illustrating the dataset composition, with 13,014 total blood samples stratified by biofluid type (serum: N=6,792; plasma: N=6,222) and diagnostic categories (control, benign, and cancer subtypes). **(c)** Schematic representation of Exai-1’s transformer-based architecture. cfRNA features, along with special tokens for contextual learning (cancer detection; CD, tissue of origin; TOO, assay version; AV, and biofluid type; BT), are embedded into a high-dimensional space. A variational encoder applies self-attention to generate a latent representation, which is then used by the decoder through generative sampling to reconstruct masked cfRNA features. Task-specific multi-layer perceptrons (MLPs) further refine embeddings for downstream applications.

Next, we assembled a large cohort consisting of 13,014 blood samples (6,792 serum and 6,222 plasma) for training and evaluating Exai-1 (Figure 1b). During Exai-1’s pre-training, we selected 7,349 cfRNA features and applied masked language modeling, by randomly masking 25% of the features among all samples of the mini-batch to encourage contextual learning (Figure 1c). To incorporate sequence-derived information, we initialized the embedding layer parameters using principal components of cfRNA sequences from RNA-FM (Chen et al., 2022) (see Methods). Each 32-dimensional sequence-based embedding was then scaled by the corresponding cfRNA abundance in each sample, ensuring that both sequence context and expression levels were captured in a single, unified representation. These embeddings were fed into Exai-1’s transformer encoder, which projects them onto a lower-dimensional latent space, where self-attention mechanisms model dependencies across cfRNA species. A variational layer then learns a latent Gaussian distribution, from which the transformer decoder samples to reconstruct the original cfRNA abundances.

Another critical design aspect that enables Exai-1 to learn sample-specific representations is the integration of special tokens for multi-task learning. These include tokens for ‘cancer detection’ <CD>, ‘tissue-of-origin’ <TOO>, ‘assay version’ <AV>, and sample ‘biofluid type’ <BT> (Figure 1c). These special tokens are essential for enhancing the model’s ability to incorporate additional contextual information alongside cfRNA abundances, enabling the model to generate biologically and clinically relevant embeddings. Consequently, the inclusion of special tokens significantly broadens the applicability and utility of Exai-1. Moreover, by applying self-attention to a compressed hidden space rather than the original high-dimensional input features, Exai-1 maintains a compact architecture with only 3.6 million trainable parameters, thus minimizing the risk of overfitting.

### Exai-1 enables high-fidelity cfRNA reconstruction and noise removal

As a masked language model, Exai-1 is trained primarily using a self-supervised objective to learn cfRNA abundances and their relationships. To evaluate its reconstruction performance, we conducted a test by randomly masking 25% of cfRNA features in held-out samples and comparing Exai-1’s reconstructed values to the original measurements. In genomic datasets, the dataset average often serves as a challenging baseline that many models struggle to surpass (Kedzierska et al., 2023; Schreiber et al., 2020). While this baseline yielded an *R*^2^ of 0.57 (95% CI: 0.55–0.59), Exai-1 significantly outperformed it, achieving an *R*^2^ of 0.89 (95% CI: 0.88–0.89, Figure 2a–b). This result underscores Exai-1’s ability to accurately capture sample-specific cfRNA abundance profiles in blood and its capability to effectively fill-in the gaps.

**Figure 2:**
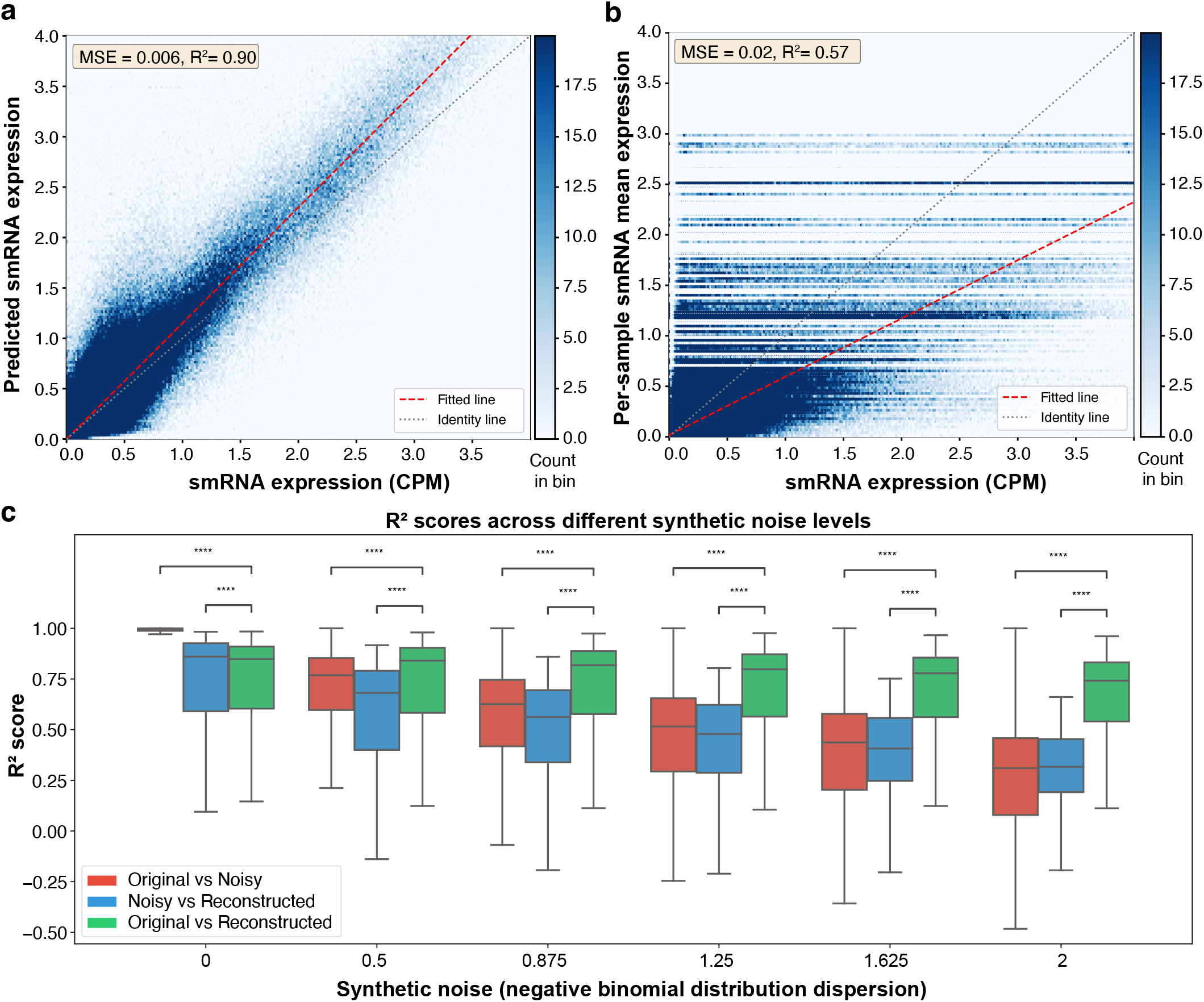
Exai-1’s performance in reconstructing and denoising cfRNA abundance data. **(a)** Density scatterplot showing Exai-1’s reconstruction of masked cfRNA features in the test set. The x-axis indicates per-sample smRNA expression levels, while the y-axis represents the reconstructed abundances. A red linear regression line illustrates the correlation. **(b)** Density scatterplot comparing the original cfRNA abundances to their mean values across samples (baseline). **(c)** Box plots summarizing Exai-1’s denoising capability across six simulated “noisy” datasets, each generated with increasing dispersion from a negative binomial distribution. Shown are *R*^2^ values comparing the noisy data and Exai-1’s reconstructions against the original, showcasing Exai-1’s robust performance in the presence of noise.

We performed ablation analyses to better understand the contribution of each component of Exai-1. Lack of self-attention, for example, significantly dropped the performance of classification tokens. On reconstruction tasks, we particularly observed a significant drop when more than 50% of the held-out dataset tokens were masked. Similarly, a model without the variational component (vanilla transformer instead of a variational transformer), achieved the lowest performance in both reconstruction and token-specific tasks (Supplementary Figure 2).

We hypothesized that reconstruction capabilities of Exai-1 should also allow us to increase the signal to noise ratio of liquid biopsy datasets. To assess our hypothesis, we introduced controlled noise into the original cfRNA expression profiles. Specifically, we modeled the observed cfRNA abundances using a negative binomial distribution and systematically increased the dispersion parameter to generate six datasets representing varying levels of noise in data. For each noise setting, we compared how closely both the noisy inputs and Exai-1’s reconstructions matched the original, noise-free abundance profiles. Exai-1 consistently produced reconstructions that were substantially closer to the original data than the noisy inputs themselves (Figure 2c). Furthermore, while the *R*^2^ between noisy and original data declined monotonically with increasing noise, Exai-1’s reconstructions exhibited only a minimal decrease in *R*^2^. These findings demonstrate Exai-1’s ability not only to accurately reconstruct cfRNA profiles but also effectively denoise them, preserving essential biological signals even in the presence of substantial noise.

Building upon Exai-1’s denoising capability, we hypothesized that reconstructed features could be used to generate synthetic cfRNA profiles, thereby augmenting training datasets and enhancing cancer detection performance using standard classifiers. We had previously shown that such synthetic profiles can improve model training and generalization (Karimzadeh et al., 2024). To evaluate this, we sub-sampled our training set to smaller datasets ranging from 50 to 400 samples (doubling the sample size at each step), each repeated 10 times (Figure 3a). For each subset, we trained an XGBoost classifier to distinguish cancer from control samples, comparing two approaches: training exclusively on samples with original cfRNA profiles (*original-only*) versus training on datasets augmented with synthetic cfRNA profiles generated by Exai-1’s reconstruction (*original+reconstructed*), effectively doubling the training sample size.

**Figure 3:**
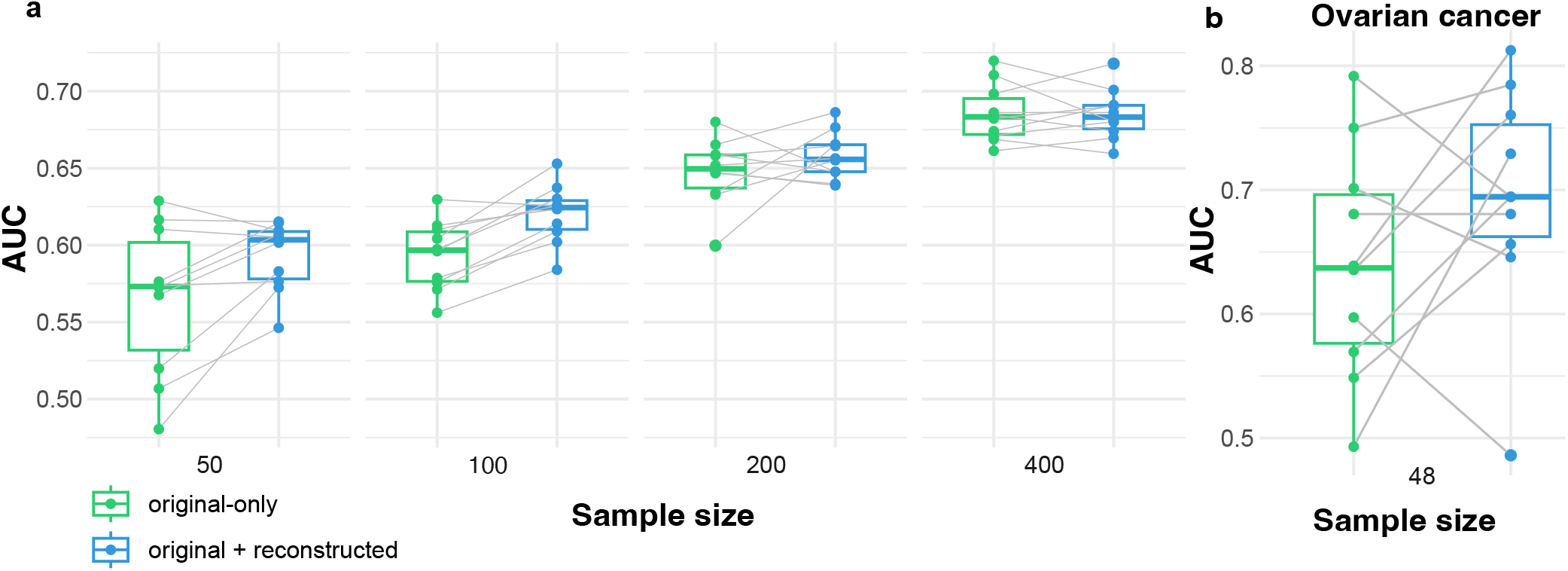
Improved cancer detection using Exai-1 synthetic cfRNA profiles. (**a**) AUC comparison for cancer detection classifiers trained either on original cfRNA samples (*original-only*) or a combination of original and Exai-1 reconstructed samples (*original+reconstructed*). Box plots summarize results across sample sizes (50–400) and 10 random seeds per size. (**b**) Classifier performance improvement for ovarian (N=48) cancer cohort when augmented with reconstructed samples. Points represent individual random seeds.

Our results demonstrate that augmenting datasets with synthetic cfRNA profiles consistently improved cancer detection performance, particularly at smaller sample sizes (up to 400 samples). A linear mixed-effects model further validated this improvement, estimating an average AUC increase of 0.032 (95% CI: 0.014–0.051, *p <* 0.005) when synthetic cfRNA profiles were included compared to using original samples alone. Such foundation-model-driven augmentation is particularly beneficial in clinical contexts where sample availability is often limited. To further test the practical utility of synthetic cfRNA profile augmentation, we applied Exai-1 to a smaller cohort of ovarian (N=48) cancer samples, supplemented with control samples from a matched study. Across ten random seeds, we observed consistent AUC improvements of 0.063 (95% CI: 0.004–0.11) for ovarian cancer detection when incorporating reconstructed data(Figure 3b). Together, these findings demonstrate the utility of Exai-1’s data denoising and synthetic cfRNA generation for enhancing cancer detection, especially in rare diseases where samples are limited.

### Exai-1’s small RNA and sample embeddings capture key biological and technical factors

Transcription, processing, and secretion of small RNA (smRNA)s are controlled by a suite of regulatory programs and pathways (Zirak et al., 2023; Garcia-Martin et al., 2022; Tosar et al., 2015). Therefore, similar to their common long RNA counterparts, smRNAs show patterns of co-expression across samples. As Exai-1 learns an embedding for each cfRNA species, we anticipate that various dimensions of this embedding will capture such co-expression blocks. To explore this possibility, we first clustered cfRNAs based on their abundance profile within our dataset. For each cluster, we then generated a representative embedding by averaging the embeddings of individual cfRNAs across each dimension. We partitioned the features into 10 abundance-based clusters of small RNAs and then assessed whether each cfRNA embedding dimension was significantly associated with any of these clusters by randomly shuffling cluster memberships and recalculating cluster embeddings. For each dimension, we compared these randomized values against the real values to construct a null distribution. This analysis revealed that smRNA co-expression clusters were significantly associated with distinct cfRNA embedding dimensions (FDR *<* 1%; Figure 4a). Each embedding dimension was enriched in an average of 2.4 (*±* 1.3 S.D.) co-expression clusters, and conversely, each cluster was associated with average of 8 (*±* 5.2 S.D.) embedding dimensions. Overall, these findings indicate that Exai-1 successfully learns sparse representations that are associated with the underlying biological characteristics of co-expressed smRNAs.

**Figure 4:**
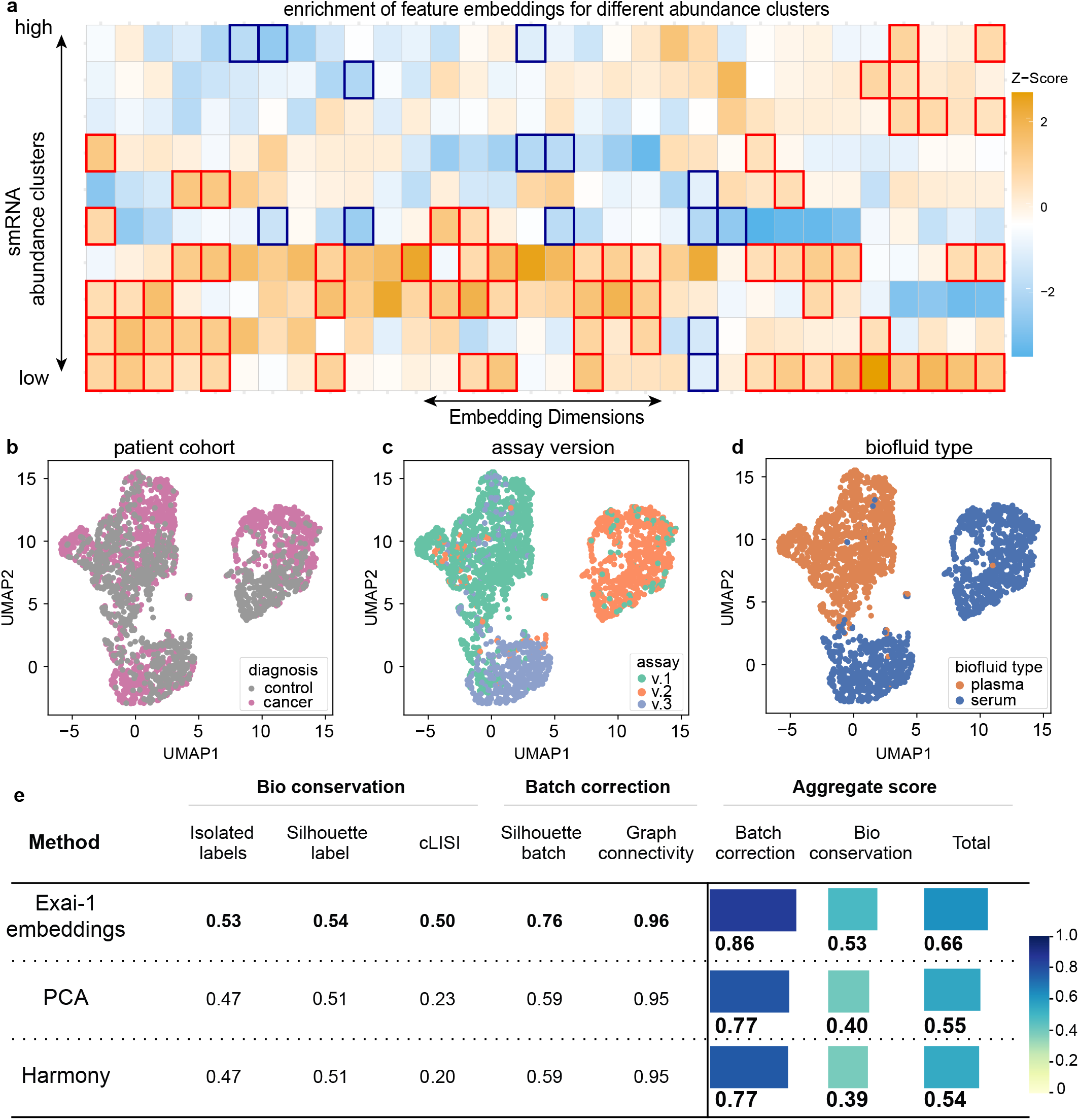
Representation of smRNA characteristics and cancer biology on feature and sample embeddings learned by Exai-1. **(a)** A heatmap depicting the standardized values of the learned feature embeddings across various abundance clusters. The significantly enriched embedding dimensions are highlighted with bold red boxes for positive and bold blue boxes for negative values. **(b)** A UMAP representation of Exai-1’s sample embeddings from the test set, color-coded by cancer and control diagnosis. **(c)** Exai-1’s sample embeddings from the test set, color-coded by assay versions. **(d)** Exai-1’s sample embeddings from the test set, color-coded by biofluid type. **(e)** The batch effect removal and biological variation conservation benchmark for Exai-1, Harmony, and PCA. Scores range from 0 to 1, with higher values indicating better performance.

Next, to visualize the 32-dimensional sample embeddings learned by Exai-1, we used UMAP to project the embedding representations onto the held-out test set. Although UMAP is designed for summary visualization and is not suitable for making inferences about the underlying data, our UMAP plots (Figure 4b–d) still reflect the observed patterns in the embedding space and highlight distinct separations between cancer and control samples, as well as differentiation based on technical variables such as assay version and biofluid type used to collect the blood samples. These visualizations indicate that Exai-1 embeddings effectively capture both biological differences and technical variations among samples.

We next assessed the impact of batch effects and the conservation of biological variance within Exai-1 embeddings, comparing them to embeddings obtained via Principal Component Analysis (PCA) and the PCA-based batch-correction method Harmony (Korsunsky et al., 2019) (Figure 4e). Following a strategy similar to (Luecken et al., 2022), we employed *k*-nearest-neighbor (*k*-NN) graph connectivity and the average silhouette width (ASW) (Büttner et al., 2019) to evaluate batch effect removal based on sample collection source. Graph connectivity measures the average proportion of samples within each cancer type connected through a *k*-NN graph, while the silhouette batch score—calculated by averaging batch-adjusted silhouette widths across cancer types—quantifies batch mixing quality (values closer to 1 indicate optimal batch integration). To assess conservation of biological variance, we used label conservation metrics, including cell-type local inverse Simpson index (cLISI) (Korsunsky et al., 2019), cancer-type silhouette width, and ‘isolated label scores’ (Luecken et al., 2022).

During training, Exai-1 leverages triplet margin loss to minimize the distance of the samples from the Gaussian latent distribution which share a common tissue of origin and are distant (see methods). Due to this label-agnostic batch correction, Exai-1’s embeddings significantly outperformed both PCA and Harmony across all metrics related to batch effect removal and conservation of biological variance. This result can be attributed to the biologically-informed contrastive loss in shaping the Exai-1’s sample and feature embeddings, leading to the learning of a biologically rich embedding space. Collectively, this result demonstrates that Exai-1 effectively captures and reflects the biological patterns in the learned embedding spaces, facilitating numerous downstream applications, especially in disease detection and diagnostics, even in limited-sample scenarios leveraging few-shot learning.

### Leveraging pre-trained Exai-1 to enhance generalizability across sample source

Although serum is more widely available in many clinical settings, plasma samples typically provide higher-quality cfRNA data with lower background noise and are therefore a more favorable biofluid for development of liquid biopsy assays. This disparity in both availability and quality poses a significant challenge in developing robust liquid biopsy classifiers, as classifiers trained solely on one biofluid type often fail to generalize to another. We therefore investigated whether the foundational capabilities of Exai-1 could allow training of more generalizable classifiers for tasks such as detection of cancer.

To evaluate this hypothesis, we used paired plasma and serum samples from the same patients which were not used during Exai-1 training. First, we used the original 7,349 smRNA log1p CPM values to train a cancer detection XGBoost model using 391 plasma samples from individuals with cancer and 485 from individuals without cancer. The cross-validated performance of this model on plasma samples achieved AUROC of 0.74 (95% C.I. 0.7–0.77) while on serum samples, this model only had an AUROC of 0.56 (95% C.I. 0.51–0.61, Figure 5a). In a similar setup, however, when we used the latent space of Exai-1 for training a plasma-specific XGBoost model, the model had a higher and comparable performance on both plasma and serum (Figure 5b).

**Figure 5:**
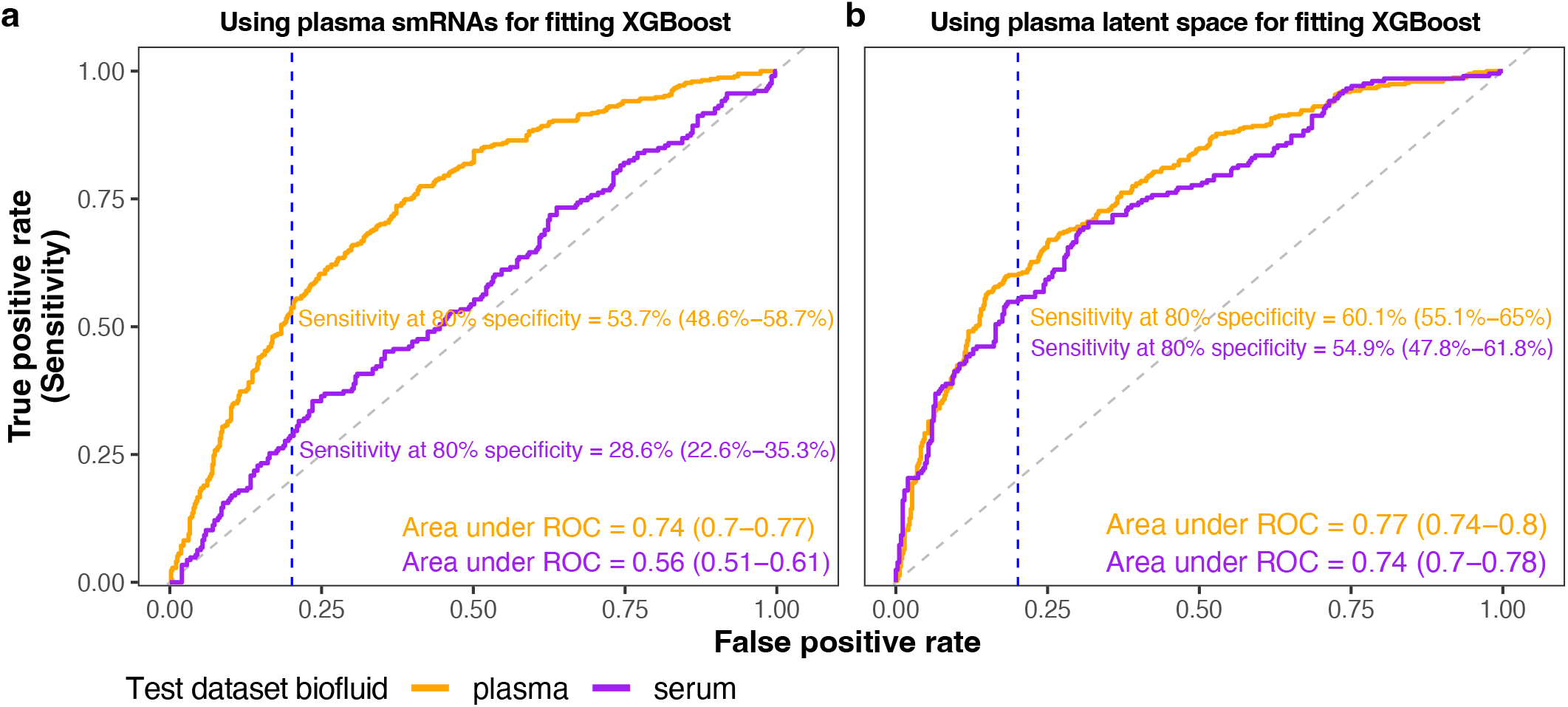
Exai-1 allows generalizability for out-of-distribution datasets. **(a)** ROC plot shows performance of an XGBoost model trained using 5-fold cross validation on plasma smRNAs. Orange shows cross-validated performance of the model on plasma samples, while purple shows performance of the model on serum samples. **(b)** Similar to **(a)** with the key difference of using Exai-1’s plasma sample latent space for training and evaluating XGBoost models.

These results highlight Exai-1’s potential to effectively address data-type biases inherent in liquid biopsies, making biofluid-specific classifiers broadly applicable to out-of-distribution datasets and thus enhancing the robustness and practical utility of future predictive tools in liquid biopsy-based cancer diagnostics.

## Discussion

In this study, we introduced Exai-1, a multi-modal generative model pre-trained on a large dataset of cell-free small RNA (smRNA) profiles derived from over 13,000 plasma and serum samples. By integrating RNA-FM–based sequence embeddings with cfRNA abundance data, Exai-1 learns rich representations that capture both the structural properties of small RNAs and the patterns in their abundance. This multi-modal approach allows Exai-1 to bridge the gap between RNA sequence context and cell-type-specific expression dynamics, enabling more biologically meaningful learning.

Exai-1 demonstrates robust performance across clinically relevant tasks, highlighting its potential as a transformative platform for liquid biopsy applications. Its generative capabilities allow for the accurate reconstruction of masked cfRNA profiles, capturing both biological variation and technical artifacts present in cfRNA data. This property facilitates effective data denoising and enables the generation of synthetic cfRNA profiles, offering a practical solution for augmenting training datasets, particularly in settings where clinical sample availability is limited. Furthermore, Exai-1 embeddings inherently reflect known biological co-expression patterns, preserving clinically relevant structure in a low-dimensional latent space. By capturing biologically meaningful variance while minimizing technical batch effects, Exai-1 emerges as a robust foundation model suitable for a range of clinical diagnostic applications.

One of the major challenges addressed by Exai-1 is the variation introduced by different biofluid types, such as serum versus plasma samples, which often limits the generalizability of conventional classifiers. Serum and plasma each have their own advantages and disadvantages with respect to biobank availability and biomarker signal to noise ratio. Exai-1 decouples biological signal from technical variations within the latent space, making it possible for a plasma-trained model to generalize to out-of-distribution serum data, a task which was not possible in the original feature space (Figure 5a). This was largely due to the novel approach we used for minimizing unknown technical variations through triplet margin loss, as well as the sparsity constraints within Exai-1’s variational inference objectives. This feature is particularly important for improving diagnostic accuracy and expanding the clinical applicability of cfRNA-based models across diverse patient populations and clinical contexts.

Previously, we showed the power of variational inference for modeling cancer-specific oncRNAs where we outperformed conventional methods such as XGBoost (Karimzadeh et al., 2024). Enriching for a cancer-specific feature space, as we showed, allows for training more accurate cancer-specific classifiers. In line with previous observations, we still observe that when using identical samples, the latent space learned through Exai-1 allows for training better classifiers than when using the original feature space Figure 5. This further confirms that Exai-1’s representation learning enhances biological signals while disentangling them from confounders, leading to more generalizable classifiers.

Unlike models trained on textual data such as ChatGPT, which benefit from vast amounts of labeled data, Exai-1 operates in a domain where ground truth samples are inherently scarce and expensive to generate. While NLP foundation models improve as their datasets expand (Zhou et al., 2023; Jaegle et al., 2021), cfRNA-based models must navigate data constraints that limit the direct scalability of their training sets. However, as larger and more diverse cfRNA datasets become available, we anticipate continued improvements in Exai-1’s performance and clinical utility.

Future development efforts will focus on enhancing Exai-1 by expanding its training datasets, incorporating additional multi-modal inputs, and refining its contextual special tokens to further improve disease classification and biomarker discovery. These improvements will position Exai-1 as a versatile foundational model, capable of significantly advancing non-invasive cancer diagnostics. In particular, Exai-1 is well-suited for challenging applications such as rare cancer detection and minimal residual disease monitoring, where data limitations have traditionally constrained model performance.

## Methods

### Dataset

Here, we utilized an in-house dataset of serum and plasma collected from 13,014 samples sourced from 11 different suppliers and more than 128 different collection sites. These included 7,265 samples from individuals with cancer and 5,749 samples from individuals without known history of cancer. We used RNA isolated from 0.5–1 mL of serum or plasma from each donor to generate and sequence cfRNA libraries for each sample, achieving an average depth of 46 *±* 26 million single-end reads. Tumor of the patients with cancer originated from breast (2,956), lung (1,932), colorectum (753), kidney (313), stomach (307), pancreas (213), bladder (169), prostate (156), liver (145), uterus (95), esophagus (91), ovary (71), and cervix (64). We used 8,339 samples for training, 2,079 for hyperparameter tuning and model selection (validation set), and 2,596 samples for reporting the performance metrics throughout the manuscript.

### Feature Selection

We curated a feature set comprising 799,289 small RNAs. Our selection process focused on tRNA, snoRNA, miRNA, yRNA, lncRNA, and oncRNAs. The library of oncRNAs were discovered through de-novo loci forming (Fish et al., 2018; Karimzadeh et al., 2024) and association analyses comparing small RNA-sequencing between tumor and adjacent normal tissue samples in The Cancer Genome Atlas (TCGA). We first excluded any feature detected in fewer than 2% of samples, thereby removing exceedingly rare cfRNAs. We next imposed a coefficient-of-variation (CV) threshold of 1, retaining only those features exhibiting sufficient variability across the data (Supplementary Figure 1a–e).

This step resulted in 394,175 oncRNAs, 716 miRNAs, 704 tRNAs, 761 yRNAs, and 610 snoRNAs. To refine the oncRNA subset further based on sequence and structure information, we used RNA-FM (Shen et al., 2024) embeddings, extracting the 32 principal components (PCs) that collectively captured over 91% of the variation. Based on these PCs, we applied an iterative k-means clustering strategy: starting with 1,000 random clusters. Each cluster with more than 10 members was iteratively divided accordingly to result in 10 or less oncRNAs. The oncRNA closest to the centroid was chosen as single representative per subcluster. This process yielded 4,558 oncRNAs for inclusion in subsequent analyses (Supplementary Figure 1e–f).

### Architecture of Exai-1

Exai-1 is inspired by the success of variational auto-encoders in modeling sparse genomic datasets (Lopez et al., 2018), masked-language modeling applications in genomics (Cui et al., 2024; de Lima Camillo et al., 2024), power of leveraging other foundation models for token embedding (Rosen et al., 2023), and capability of combined transformer and variational Bayes architectures in capturing high-level context (Wang and Wan, 2019). Exai-1 facilitates the handling of large-scale cfRNA data without over-parameterizing and overfitting to training data.

Let 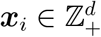 denote log1p of counts per million (CPM) mapped reads for *d* − 4 smRNAs of the *i*-th sample. We appended 4 additional rows, ***x***_*i,d*−3_, …, ***x***_*i,d*_ as placeholders for classification tokens (cancer detection and tissue of origin) as well as sample information tokens (collection tube type and assay version). While classification tokens are always masked by setting them to -1, sample information tokens are always provided. Since we have four classification and sample information tokens, each with *t*_*b*_ different targets, *b* = 1, …, 4, these tokens can be represented as 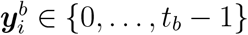.

For encoding each smRNA token, we leverage the sequence and structure-aware embeddings of the RNA-FM model by initializing the weights of the embedding layer with a min-max-scaled version of the principal components of RNA-FM embeddings, ensuring range of weights to be within [−0.1, 0.1] for numeric stability.

Exai-1 encoder works as follows:

1. Token embedding step transforms small RNAs (χ) into embedding space 𝒢 (*f*_*t*_ : χ→ 𝒢). As a result, vector 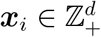 will be projected into embedding matrix 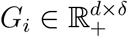 with the embedding dimension δ = 32.
2. Signal embedding step performs element-wise multiplication of smRNA embeddings with log1p CPM level of expression, resulting in *f*_*g*_ : 𝒢→ ℛ.
3. A dimensionality reduction layer with 2-dimensional layer normalization to prevent any numeric instability and a multi-layer perceptron to reduce the dimensions from *d* to 128: *f*_*h*_ : ℛ →ℋ.
4. A self-attention transformer layer: 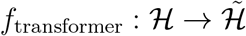.
5. A second dimensionality reduction layer generating parameters of a diagonal Gaussian distribution for variational inference, **µ**_*i*_ and **σ**_*i*_.

The self-attention layer computes attention scores over the reduced dimension of the hidden layer and embedding dimension δ = 32 through generation of:

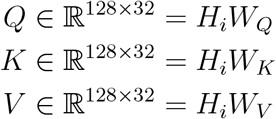

In Exai-1, the attention output 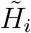, therefore, is a results of 2 layers of:

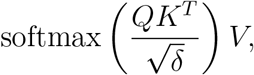

each followed by a 2-layer of dense layers with dimensions of 128, GELU activation layer (Hendrycks and Gimpel, 2016), layer normalization, and a residual connection involving bilinear interpolation of .30*R*_*i*_. This step allows each feature representation within *H*_*i*_ to attend to other variables within the same reduced feature space, thereby enhancing the model’s ability to capture intricate, context-aware representations of the latent variables.

Exai-1 attempts to learn a probabilistic low-dimensional latent space ***z***_*i*_ from 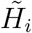, ensuring preservation of biological information. To achieve this, Exai-1 flattens 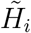 into 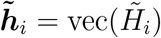 and uses a dense layer to learn the parameters of the diagonal Gaussian distribution ***z***_*i*_ ∼ 𝒩 (***µ***_*i*_, ***σ***_*i*_*I*).

Exai-1’s decoder, therefore, samples from ***z***_*i*_ during training to reconstruct the original input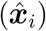. Similar to the encoder, Exai-1’s decoder involves:

1. A dense layer to increase the dimension from ***z***_*i*_ to 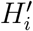
2. A self-attention decoder 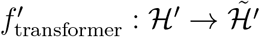 with identical architecture as *f*_transformer_ but with independent parameters
3. A dense layer to map the dimensions to the original embedding space: 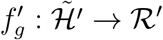
4. MLPs with 20% dropout, a hidden layer of dimension 16, and a LeakyRELU activation function. These MLPs are responsible for smRNA expression reconstruction as well as token classification.

To emphasize the preservation of biological information during dimension reduction of Exai-1, we developed a novel approach for selection of positive and negative anchors for computation of the triplet margin loss. Previously, we showed that through explicit definition of known technical variables (e.g., batch effects), we can minimize solely technical differences among the samples through triplet margin loss (Karimzadeh et al., 2024). Here, we show that without any knowledge of technical sources of variation, we can remove batch effects while preserving biological information. We achieved this through the definition of known biological information of the samples during training (e.g., tissue of origin for cancer samples).

For each sample, we identify all possible “positive” anchors 𝒥, *i* ∉ 𝒥, such that they share the same classification label ***y***_*i*_ = ***y***_*j*_ for all *j* ∈ 𝒥. We compute the Euclidean distance (*d*) among all pairs of ***z***_*i*_ and ***z***_*j*_ and sample positive anchors according to the weights set through:

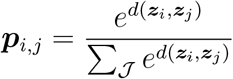

Samples of same biological label that are more distant in the embedding space, therefore, will become closer. For sampling negative anchors, we randomly sample from the pool of all samples with a different biological label.

#### 1. Reconstruction through masked tokens

During training, we randomly mask 25% of small RNAs of ***x***_*i,d*−4_ at each mini-batch among all samples, including non-expressed small RNAs, generating ***x***_*M*_, akin to the masked language modeling used in BERT (Devlin et al., 2019) and other similar models. The reconstruction Smooth L1 loss (Girshick, 2015), therefore, can be represented as:

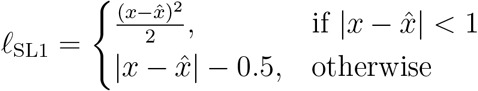

We compute this loss once for *x* and once for ***x***_*M*_.

#### 2. Kullback Leibler divergence

We minimize

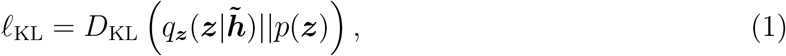

where *D*_KL_ is the Kullback-Leibler (KL) divergence (Kullback and Leibler, 1951) and *p*(***z***) = 𝒩 (**0**, *I*) is the prior distribution for ***z***, a standard Gaussian.

#### 3. Triplet Margin Loss

For each sample *i*, we sample *ω* triplets as follows:

a. Randomly pick a “positive” anchor *j* ≠ *i* such that they share the same classification label ***y***_*i*_ = ***y***_*j*_ and the sampling probability is assigned according to the distance of the two samples within ***z***_*i*_.
b. Randomly pick a “negative” anchor *j*^*′*^ ≠ *i* such that they do not share the same classification label ***y***_*i*_ ≠ ***y***_*j*_^*′*^.

During training we add a cost function that minimizes the differences among the samples that share the same biological label (e.g., cancers of the same tissue), while moving samples with different labels (e.g., cancer samples from non-cancer samples) further apart:

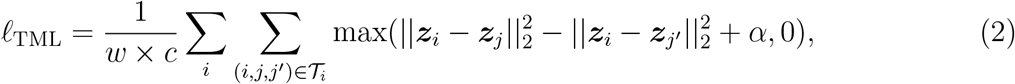

where α is a hyperparameter that enforces what should be the minimum difference of distances between a sample and its positive and negative anchors in the latent space, and it is set to *α* = 0.1.

#### 4. Cancer detection

We require token *d* − 1, which is always initialized as missing value indicator -1, to predict presence of cancer by minimizing the cross entropy loss:

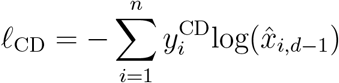

#### 5. Tissue of origin

We also require token *d*, which is always initialized as missing value indicator -1, to predict tissue of origin in presence of cancer through convergence of an additional cross entropy loss:

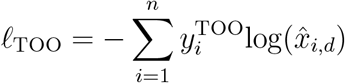

#### 6. Assay version

We require token *d* − 2, which is always according to the assay version of the sample of interest, to preserve this information in the decoded output through convergence of an additional cross entropy loss:

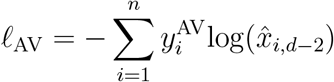

#### 7. Biofluid type

We also require token *d* − 3, which is always according to the biofluid (serum or plasma), to preserve this information in the decoded output through convergence of an additional cross entropy loss:

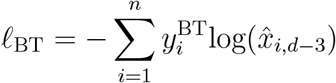

Therefore, Exai-1 attempts to converge the following loss function:

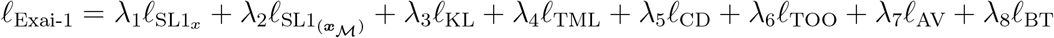

Hyperparameters for training Exai-1 were set through grid search and the selected combinations included *λ*_1_, *λ*_2_ = 100, *λ*_3_, *λ*_5_ = 10, *λ*_4_, *λ*_6_, *λ*_7_, *λ*_8_ = 1, learning rate of 0.0001, weight decay of 0.00001, gradient clipping value of 0.1, mini batch size of 256, and training for 5,000 epochs. The efficient architecture of Exai-1 allowed us to train the model over 1 NVIDIA T4 GPU for 60 hours.

## Acknowledgments

The results shown here are in part based upon data generated by the TCGA Research Network: https://www.cancer.gov/tcga. HG is a core investigator at Arc Institute. This study is funded by Exai Bio Inc.

## Supplementary Information

**Supplementary Figure 1:**
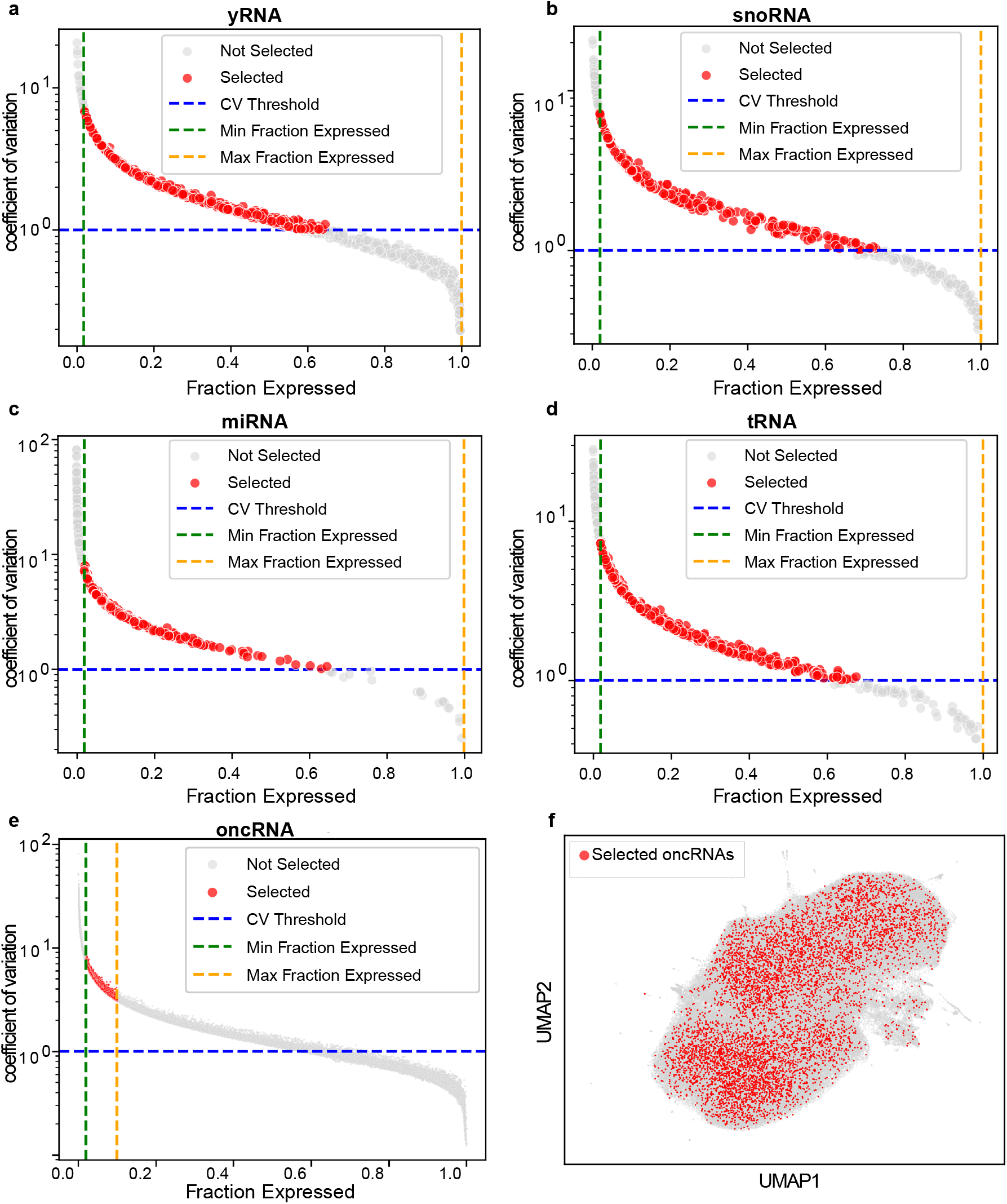
Feature selection workflow for Exai-1. **(a)** X-axis shows the fraction of samples expressing each yRNA and Y-axis shows the coefficient of variation (CV). We used the subset of samples that surpassed the CV threshold (blue dashed line) while falling within minimum and maximum fraction-expressed cutoffs (green and orange dashed lines, respectively). **(b–e)** similar to **(a)** for snoRNA, miRNA, tRNA, and oncRNA, respectively. **(f)** UMAP projection of RNA-FM embeddings for the initially retained cfRNAs, highlighting in red the 4,558 representative oncRNAs selected from an initial pool of 394,175.

**Supplementary Figure 2:**
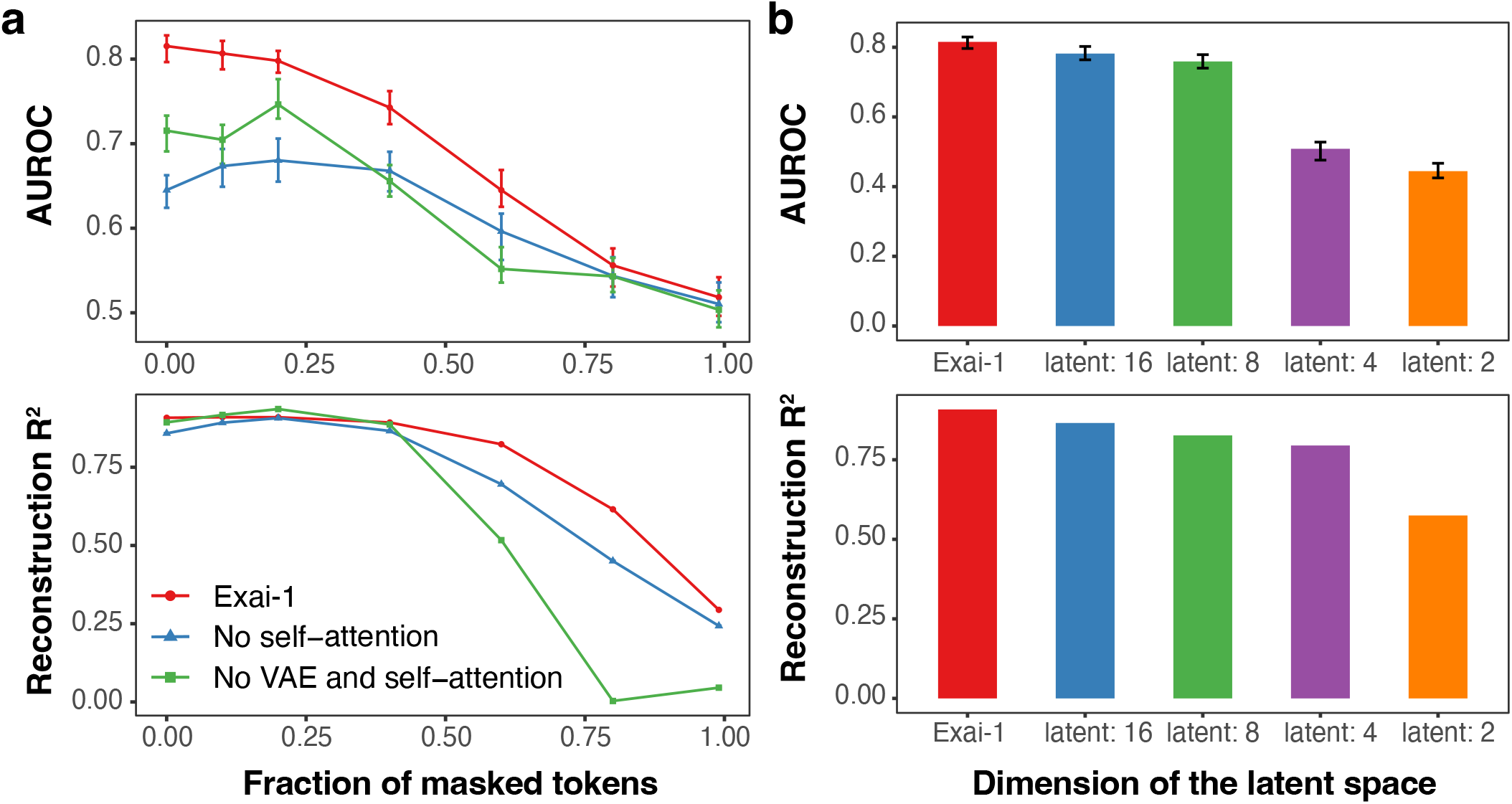
Hyperparameter impacts on Exai-1 performance. **(a)** We examined the performance of Exai-1 (red) compared to an ablated Exai-1 model without the transformer components (blue) or without the entire transformer-VAE component. Top panel shows the AUROC for cancer detection task (y-axis) when masking different percent of tokens (x-axis). Similarly, bottom panel shows the reconstruction performance as measured by *R*^2^ (y-axis). **(b)** We examined the impact of different latent dimensions on AUROC of the cancer detection token (top) as well as the reconstruction task *R*^2^. X-axis shows Exai-1 (red) with optimal latent size of 32 and other models using a smaller latent size (2––16).

